# HXMS: a standardized file format for HX-MS data

**DOI:** 10.1101/2025.10.14.682397

**Authors:** Kyle C. Weber, Chenlin Lu, Roberto Vera Alvarez, Bruce D. Pascal, Anum Glasgow

## Abstract

**Motivation:** Hydrogen/deuterium exchange-mass spectrometry (HX-MS) is a rapidly expanding technique used to investigate protein conformational ensembles. The growing popularity and utility of HX-MS has driven the development of diverse instrumentation and software, resulting in inconsistent, non-standardized data analysis and representation. Most HX-MS data formats also employ only mean deuteration representations of the data rather than full isotopic mass spectra, which reduces the information content of the data and limits downstream quantitative analysis.

**Results:** Inspired by reliable protein structure and genomics data formats, we present HXMS, a unified, lightweight, scalable, and human-readable file format for HX-MS data. The HXMS format preserves the isotopic mass envelopes for all peptides, captures the full experimental time-course including fully deuterated control samples, and contains all other key information. It supports multimodal distributions, post-translational modifications (PTMs), and experimental replicates. To promote compatibility with existing HX-MS workflows, we also developed PFLink, a Python package that converts exported data files from commonly used HX-MS software to the HXMS format. PFLink and the HXMS format will enable more quantitative, higher-resolution data processing, improved data sharing and storage among HX-MS practitioners, future machine learning applications, and further developments in HX-MS analysis.

**Availability and implementation:** PFLink is publicly available to install locally on HuggingFace, alongside documentation, or use online at HuggingFace (https://huggingface.co/spaces/glasgow-lab/PFlink). The supplemental information includes sample input files, sample HXMS files, and a generic unfilled PFlink custom CSV file that users may populate with key experimental conditions and results, which can then be read and converted into the HXMS format.

## 1 Introduction

Hydrogen/deuterium exchange-mass spectrometry (HX-MS) is a powerful biophysical method for probing protein folding and conformational ensembles (Englander and Kallenbach 1983; Englander *et al*. 2016). HX-MS measures the rate at which backbone amide hydrogens in a protein are replaced with deuterium atoms when exposed to deuterated buffers, providing time-resolved insights into protein structure and conformational stability (Hamuro 2024). The availability of automated systems to perform HX-MS experiments (Wales *et al*. 2008; Chalmers *et al*. 2006a; Wei *et al*. 2014; Chalmers *et al*. 2006b; Kish *et al*. 2023) and analyze HX-MS data has increased its appeal for studying large, complex proteins, allowing scientists to probe many pharmacological targets for applications such as drug discovery, epitope mapping, and protein characterization (Gertsman *et al*. 2009; Wang *et al*. 2012; Hamuro 2017; Gramlich *et al*. 2021; Glasgow *et al*. 2023; Jia *et al*. 2023; Shaw *et al*. 2023; Wales *et al*. 2024; Wells *et al*. 2025).

Despite the widespread adoption of HX-MS in the structural biology community, most practitioners analyze data at the peptide level using the mean deuteration of the isotopic mass envelope at each time point. This quantity is often referred to as the “centroid” representation of the data, although this usage differs from the mass spectrometry convention of centroiding profile spectra into stick representations. Although it is commonplace, the practice of using MS mean deuteration rather than the full MS envelopes limits opportunities for quantitative treatment of HX-MS data due to the resulting information loss and degeneracy (Kan *et al*. 2019; Lu *et al*. 2026). This problem is exacerbated by the large size and instrumentation-specific formats of raw MS data files, and the varied export formats of available HX-MS software. Altogether, while there have been efforts to standardize reporting in the field (Masson *et al*. 2019), sharing HX-MS data remains difficult and cumbersome since no suitable standardized format exists.

Here we present HXMS, a lightweight and human-readable file format for HX-MS data that includes the experimental isotopic mass envelopes and all necessary information for high-resolution data analysis. We also introduce PFLink, a software package that converts HX-MS data files produced by four different commercial and academic software packages, or our custom data input file, to the HXMS format (Fig. 1). The standardized HXMS format will advance data processing, data sharing, and technical developments within and beyond the HX-MS community.

**Figure 1.**
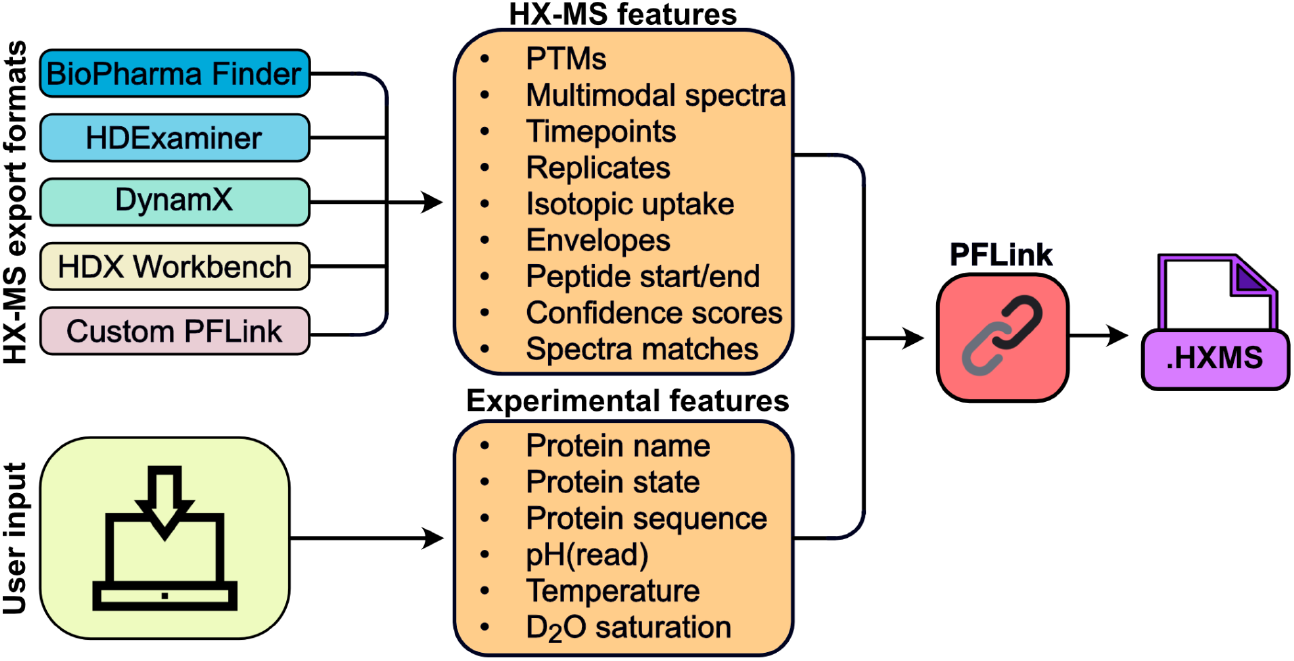
HXMS file creation using PFLink. The user provides two types of information to PFLink: 1) HX-MS data exported directly from the analysis software, or in the PFLink custom format, and 2) details about the protein and experimental conditions. PFLink automatically compiles this information in the standardized HXMS format.

## 2 Results

### 2.1 Metadata section format

The HXMS file format consists of three main sections: a metadata section, an experimental data section, and a post-translational modification (PTM) dictionary section. The metadata section describes the required experimental conditions used in the HX-MS experiment (Table 1). Important variables include the protein name, sequence, and state; the temperature, the pH(READ), and the D_2_O saturation. The user may add additional metadata using a “REMARK” header in the metadata section. Metadata objects must start with either “METADATA” or “REMARK”, followed by a tab. The title comes next, followed by a final tab before the data or remark itself, and ends with a new line character. The HXMS file version is automatically added to the “REMARK” section. The current version is v1.0.

**Table 1.** Metadata information and requirements.

| Metadata | Information | Example | Required |
| --- | --- | --- | --- |
| PROTEIN_SEQUENCE | Protein sequence | GSHMKTVEVNGADASDDN | Yes |
| PROTEIN_NAME | Name of the protein | Human PFK-1 | No |
| PROTEIN_STATE | State of the protein | APO | No |
| TEMPERATURE (K) | Temperature in Kelvin | 293.15 | Yes |
| pH(READ) | pH(read) | 6.0 | Yes |
| D2O_SATURATION | D<sub>2</sub>O saturation | 0.91 | Yes |

### 2.2 Experimental data section format

The experimental data section contains all information necessary to represent a singular timepoint in any HX-MS experiment (Table 2). In the HXMS file format, each field is represented as a column in the tabular data section. The “TP” tag is used to declare a timepoint, followed by a peptide timepoint index, “INDEX”, which incrementally increases by 1. If any timepoint represents a peptide that exhibits a multimodal distribution, the “MOD” column designates each distinct population using alphabetical indices (A–Z). A single, unimodal distribution is labeled “A” by default, whereas additional coexisting populations are assigned subsequent alphabetical indices (e.g., “B”, “C”, etc.) in the “MOD” column, thereby supporting extensible multimodal behavior. The “START” and “END” columns indicate the peptide’s beginning and end indices. These are inclusive of both ends, and indexing starts at 1 for the first amino acid in the protein. If there are multiple experimental measurements of the same peptide, the “REP” column may be indexed to denote this, where index 0 is the default for a single replicate. Replicates should be incremented only for new experimental measurements, not for timepoints in the same experiment. The “PTM_ID” column catalogs PTMs. For each unique PTM, whether it is on the same peptide or a different one, the counter is incremented. The default setting for no PTM uses the index 0000. The “PTM_ID” is used in the lookup table in the PTM section of the HXMS file to provide information about each modification, as described further in Section 2.3. The “TIME(SEC)” column indicates the duration of the sample incubation in D_2_O before it was quenched. Time is reported in scientific notation, except for fully deuterated samples, which are reported as “inf”. The “UPTAKE” column denotes the amount of deuterium incorporated. This is calculated by subtracting the mean deuteration of the 0 s timepoint from the mean deuteration for each given timepoint, for each peptide and replicate. Finally, the “ENVELOPE” column contains full isotopic mass envelopes when envelope-level spectral data are available. There is no limit to the number of peaks that can be included. The peaks are separated by commas and are normalized to sum to 1.

**Table 2.** HX-MS data.

| Header name | Information | Number of characters | Separation character | Example |
| --- | --- | --- | --- | --- |
| TITLE | Type of data | 12 | N/A | TP |
| INDEX | Peptide index | 8 | N/A | 0 |
| MOD | Multimodal index | 7 | N/A | A |
| START | Peptide start position | 7 | N/A | 1 |
| END | Peptide end position | 7 | N/A | 10 |
| REP | Experiment number | 5 | N/A | 0 |
| PTM_ID | PTM ID used for the PTM lookup table | 8 | N/A | 0000 |
| TIME(SEC) | Incubation time in D<sub>2</sub>O | 16 | N/A | 0.000000e+00 |
| UPTAKE | Mean deuteration uptake relative to 0 time point | 9 | N/A | 0.00 |
| ENVELOPE | Mass spec full envelope | inf, 4 per peak | , | 0.527,0.298,0.116,0.036,0.000,0.023,0.000 |

To account for back exchange, the HXMS format includes fully deuterated controls as standard timepoints within the experimental data section. These entries follow the same structure as other timepoints and allow replicates, with “TIME(SEC)” explicitly set to “inf” to enable consistent quantification and correction. The UPTAKE column reports raw deuterium incorporation and is not corrected for back exchange.

### 2.3 PTM section format

The PTM section serves as a dictionary, providing detailed information for any PTMs in the dataset (Table 3). This section links the ‘PTM_ID’ from the experimental data section to a comprehensive description of the modification. The ‘PTM’ tag is used to declare a PTM, followed by the ‘PTM_ID’ in the timepoint series section. The description of the entry is in the ‘CONTENT’ column, which describes the modified amino acid within the protein sequence. If there are multiple PTMs in one peptide, one may place a comma and denote the next PTM on the same line. We recommend using PDB ligand ID for simplicity and clarity, but any declaration format can be used due to the high diversity of PTMs.

**Table 3.** PTM format.

| Header name | Information | Number of characters | Separation character | Example |
| --- | --- | --- | --- | --- |
| TITLE_PTM | Type of data | 12 | N/A | PTM |
| PTM_ID | PTM ID used for the PTM lookup table | 8 | N/A | 0000 |
| CONTENT | The PTM denoted by the absolute position of the PTM in the peptide and PTM ID | inf | , | Phosphoryl STY (18) |

**Table 4.** MATCH format.

| Header name | Information | Number of characters | Separation character | Example |
| --- | --- | --- | --- | --- |
| TITLE_MATCH | Type of data | 12 | N/A | MATCH |
| TP_ID | TP ID used for MATCH lookup | 8 | N/A | 1 |
| CONF | Confidence score for the matched envelope assignment | 8 | N/A | 1.00 |
| RT(min) | Chromatographic retention time, minutes | 8 | N/A | 8.680 |
| Z | Charge state | 5 | N/A | 4 |
| MONO_M | Monoisotopic mass of the matched peptide | 16 | N/A | 2091.977598 |
| M/Z | Raw isotopic mass envelope data | inf | , separates envelope peaks corresponding to TP rows<br>; separates fine isotopologue structure within each envelope peak<br>: separates m/z and intensity | 523.9764:2.076;523.9792:2698.8441;523.982:1595.0798, 524.319:7.059;524.3218:3170.0975;524.3246:8130.5251; 524.3273:9330.6525;524.3301:7816.5479;524.3329:3718.9259;524.3357:1818.192 |

### 2.4 MATCH section format

The optional MATCH section is designed to preserve the raw matched isotopic envelope information associated with each timepoint “TP”. This section functions as a lookup table linking processed timepoint data to the original spectral evidence, thereby ensuring full traceability of peak assignments. The “MATCH” tag specifies the data type. Each entry is indexed by “TP_ID”, which corresponds to the identifier used in the timepoints. The “CONF” field records the confidence score for the matched envelope assignment. The “Z” denotes the charge state of the peptide ion. The MONO_M field reports the monoisotopic mass of the matched peptide.

The “M/Z” header stores the raw isotopic mass envelopes using a hierarchical delimiter format. Individual envelope peaks corresponding to entries in the timepoint table are separated by commas “,”. Within each envelope peak, uncentroided fine structures of the spectra, including isotopologues, are separated by semicolons “;”. For each peak or isotopologue, the *m*/*z* value and its intensity are separated by a colon “:”. This design ensures that the complete raw matched envelope information is retained in a structured and reproducible format.

### 2.5 PFLink: software to convert HX-MS files to the HXMS format

To enable widespread compatibility and accessibility with the HXMS format, we developed PFLink, a Python package to convert exported HX-MS data files from several widely used commercial and academic HX-MS data analysis programs. PFLink is compatible with exported data from BioPharma Finder (Thermo Fisher), HDExaminer (Trajan), DynamX (Waters), and HDX Workbench (Pascal *et al*. 2012) (Fig. 1, Supplemental Data 1). As most HX-MS analysis software packages currently only support the export of HX-MS data in the mean deuteration format, PFLink can write HXMS files originating from any of these programs in the mean deuteration format.

BioPharma Finder and DynamX report deuterium uptake and modifications directly, so HX-MS data exported from these programs can be used as-is. In contrast, PFLink recalculates deuterium uptake as reported by HDX Workbench by first identifying the zero timepoint for each peptide (or an average of the replicates, if any replicate is missing a zero timepoint), and then normalizes each timepoint by subtracting this zero timepoint value. PTMs are supported on all file formats except for HDExaminer and BioPharma Finder; for these programs, they must be added to the HXMS file manually.

Because both HDX Workbench and HDExaminer support exporting complete isotopic mass spectra for all peptides, PFLink can also generate HXMS files in the full-spectrum format when provided with this input. Leveraging the full isotopic mass spectra exported by HDX Workbench and HDExaminer, PFLink can generate the MATCH section, preserving either centroided envelopes or uncentroided isotopologue fine structure as structured *m*/*z*-intensity pairs, thereby maintaining direct traceability to the underlying raw spectral data.

Alternatively, users may choose to run PFLink using the custom data format supplied in Supplemental Data 1. The resulting HXMS files are compatible with the quantitative and high-resolution HX-MS analysis methods PFNet (Lu *et al*. 2025) and FEATHER (Lu *et al*. 2026).

### 2.6 Two examples

We include two sets of HXMS files on two proteins as examples (Supplemental Data 2): one set for *E. coli* DHFR in its apo state and two inhibitor-bound states (Lu *et al*. 2026), and another set for the pre- and post-fusion stabilized states of the herpes simplex virus 1 (HSV-1) glycoprotein B (gB) (Roark *et al*. 2025). In both cases, we collected the data using a Bruker MaXis II LC-QTOF mass spectrometer. Both datasets were processed using PIGEON-derived peptide lists (Lu *et al*. 2026) in HDExaminer. State-specific HXMS files were then generated using PFLink: apo, methotrexate (MTX)-bound, and trimethoprim (TMP)-bound for DHFR; and pre- and post-fusion for HSV-1 gB. The DHFR files were generated with the fine match option enabled, and include the uncentroided fine structures from the raw mass spectra. The HSV-1 gB HXMS files contain bimodal spectra. Examples of PTMs can be found in the DynamX supporting files (Supplemental Data 1).

## 3 Discussion

In this article, we introduced HXMS and PFLink: a standardized, lightweight, and human-readable HX-MS data format, and a software package to convert HX-MS data to this format. In designing the HXMS format, we drew from the most successful elements of protein structure and genomics data formats to establish a flexible framework for HX-MS data inspection, storage, and sharing. The key advantage of the HXMS data format is its inclusion of isotopic mass envelopes and all other data necessary for quantitative and high-resolution data analysis. PFLink preserves all features of the data upon conversion to HXMS: experimental and biological replicates; PTMs; multimodal distributions; all measurements, including fully deuterated samples; and, when necessary or preferable, a mean deuteration-level data representation, and/or an uncentroided data representation.

While other HX-MS data formats are widely in use, these can often only be used with specific mass spectrometers and rely on mean deuteration representations, which fail to capture the full complexity of HX-MS data. The lack of standardization in the field, combined with the large size of raw HX-MS data files, limits the extraction of valuable information from past HX-MS studies. By contrast, the HXMS format is compatible with major commercial and academic HX-MS analysis software packages, including those enabling accurate and quantitative determination of high-resolution ensemble energies from HX-MS data (Kan *et al*. 2013; Gessner *et al*. 2017; Saltzberg *et al*. 2017; Babić, Kazazić and Smith 2019; Skinner *et al*. 2019; Salmas and Borysik 2021; Smit *et al*. 2021; Stofella *et al*. 2022; Puchała *et al*. 2025; Lu *et al*. 2025; Lu *et al*. 2026).

While the HXMS format aims to be a unified format for sharing HX-MS data, it is not intended to replace information-dense raw data files, but rather to be complementary to them. We strongly recommend that practitioners upload all raw data to repositories such as ProteomeXchange (Vizcaíno *et al*. 2014; Deutsch *et al*. 2023), while including HXMS files as supplementary data files with their publications. We hope that commercial and academic vendors of HX-MS analysis software packages will onboard HXMS format export capability.

Because upstream processing of raw HX-MS data directly impacts peptide selection and isotopic mass envelope extraction, vendor-specific handling can obscure how the data were processed. We address this by adding the MATCH section to the HXMS format, which ensures transparency regarding vendor-specific biases. The full traceability of peak assignments will allow for the review of isotopic mass envelope processing for debugging and other purposes without the need for vendor-specific software, greatly increasing the accessibility of the data. The MATCH section maintains a human-readable format to further assist with the inspection and debugging process.

Overall, the HXMS format will improve data parsing, analysis, sharing, and storage for the HX-MS community and enable future machine learning and integrated structural biology applications that require large amounts of HX-MS data from many practitioners. As MS technology develops, and the needs of our HX-MS community grow and change, the HXMS data format will evolve to accommodate HX-MS theoreticians, practitioners, and software/technology developers, as well as newcomers to the field.

## Acknowledgements

We acknowledge HX-MS data collection by Andrew Reckers for DHFR and Dr. Malcolm Wells for HSV-1 gB, as well as software testing and features requests by Savannah McBride and other members of the Glasgow Lab, Dr. Andrea Piserchio, Dr. Rinat Abzalimov, and discussions with Vlad Sarpe (Trajan) and Dr. Yoshitomo Hamuro (Johnson & Johnson). This work was supported by the National Institutes of Health (R35GM157185) and a National Science Foundation Graduate Research Fellowship to KCW.

